# Deep learning discerns cancer mutation exclusivity

**DOI:** 10.1101/2020.04.09.022731

**Authors:** Prashant Gupta, Aashi Jindal, Jayadeva, Debarka Sengupta

**Affiliations:** Department of Electrical Engineering, Indian Institute of Technology Delhi, Hauz Khas, Delhi 110016, India; Center for Computational Biology, Indraprastha Institute of Information Technology, Delhi 110020, India; Department of Computer Science and Engineering, Indraprastha Institute of Information Technology, Delhi 110020, India

## Abstract

The exclusivity of a vast majority of cancer mutations remains poorly understood, despite the availability of large amounts of whole genome and exome sequencing data. In clinical settings, this markedly hinders the identification of the previously uncharacterized deleterious mutations due to the unavailability of matched normal samples. We employed state of the art deep learning algorithms for cross-exome learning of mutational embeddings and demonstrated their utility in sequence based detection of cancer-specific Single Nucleotide Variants (SNVs).

---

The widespread adoption of high-throughput DNA sequencing technologies has allowed the sequencing of the genomes/exomes of several thousands of individuals^1^. However, the sheer size of the human genome has ever bottle-necked the quest for a universal and comprehensive understanding of variations, cropping up from the sequence data deluge. We propose the *Aminoacid Switch Sequence Model* (ASSM) to represent all possible coding SNVs (Fig. 1A). ASSM encodes an SNV as a switch of aminoacids. For all 640 (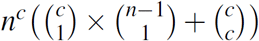 where codon length, *c* = 3, and number of nucleotides, *n* = 4) such switches, we learned continuous representation by scanning ∼ 60,000 exomes, archived by the Exome Aggregation Consortium (ExAC)^1^. Due to significant computational overhead for the current study, we initially focused on the sex chromosomes in the present study. We note that ∼ 70 protein-coding genes harbored by Chromosome Y offer inadequate levels of genetic diversity, thereby trivializing deep learning-based interventions. For Chromosome X, we obtained about 107,000 high-quality variants from the ExAC browser. We constructed Aminoacid Switch Sequence for all these alterations while accounting for the splicing events. The switch sequences were subjected to *Skipgram*^2^, a single layer neural network, widely used for learning *word-embeddings* from large text corpora. The semantic representation thus obtained was captured in a 300 sized numeric vector representing the individual switches. As the vectors of similar words correlate highly, switches that share similar nucleotide contexts orient themselves analogously in the associated vector space. We restricted the scope of the Skipgram based representation learning to the ExAC reported SNVs alone since they belong to populations that are largely healthy. In this way we conjectured, variations linked to diverse diseases can be recognized easily.

**Figure 1.**
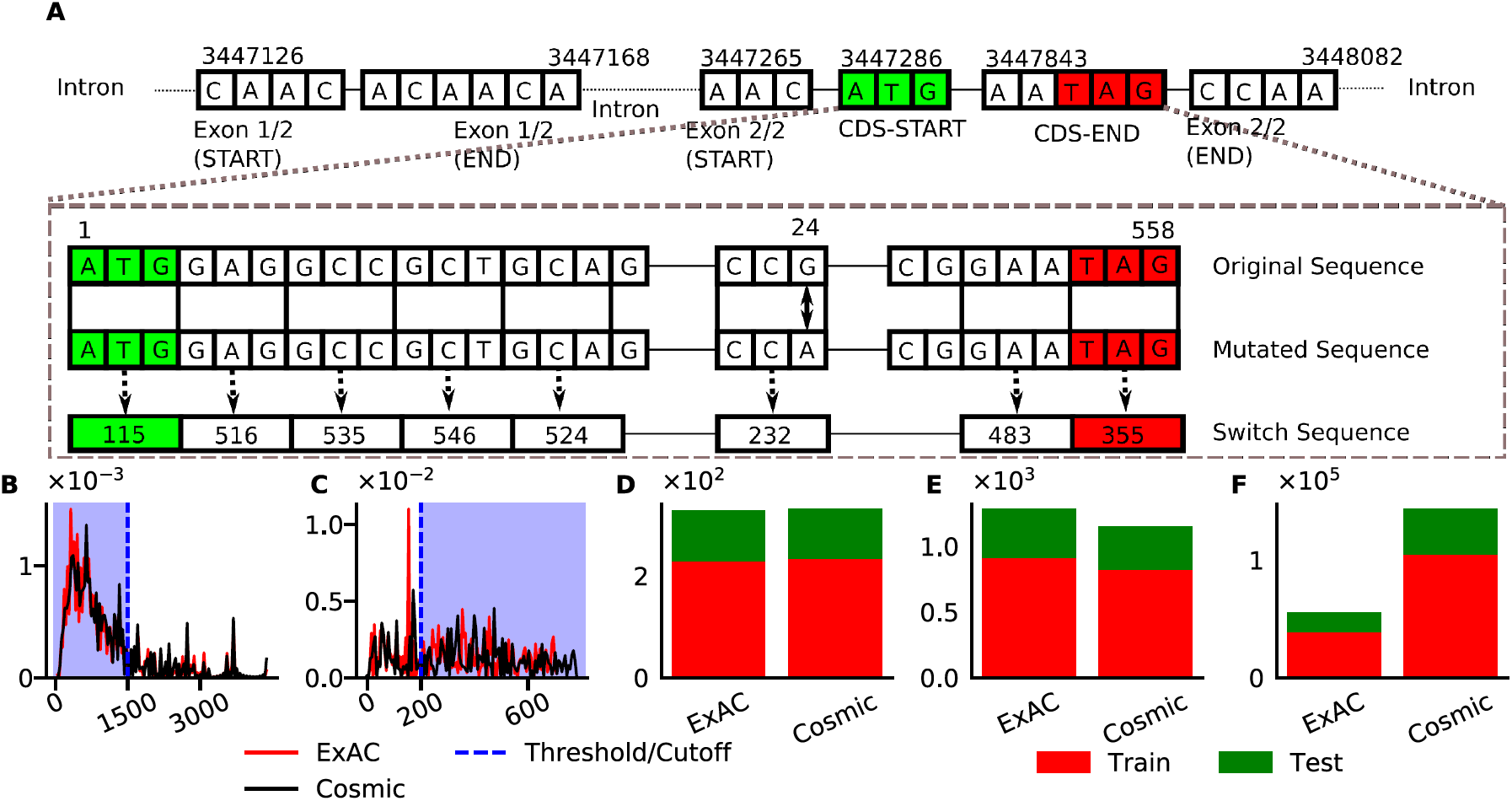
(*A*) A toy example illustrating the process of constructing Aminoacid Switch Sequences. Numerals in the switch sequence correspond to the index of switch in switch dictionary. (*B*) Distribution of the switch sequence lengths corresponding to splice variants. Sequences containing fewer than 1500 switches were used for the classification tasks. (*C*) Distribution of SNV-featuring splice variants per gene. Genes fewer than 200 such sequences were filtered out. (*D*) Barplots depict the counts of Chromosome X specific genes that contain the SNVs. These numbers are 328 and 332 for ExAC and COSMIC, respectively. (*E*) Barplots depict the splice variant counts for these genes. For ExAC and COSMIC these numbers are 1286 and 1153, respectively. (*F*) Barplots depicting the counts of Aminoacid Switch Sequences after considering the SNVs (38,754 and 105,170 for ExAC and COSMIC respectively). Note: 70/30 rule was applied on genes to construct the training and validation sets.

Cancer related somatic mutations are of two broad types – drivers and passengers. Drivers confer a fitness advantage to cancer cells, whereas passengers, *aka*. “hitchhikers,” don’t^3^. Passenger mutations comprise about 97% of somatic SNVs in cancer. Recent shreds of evidence highlight the indirect and damaging roles of passenger mutations^4^. The analysis of cancer genomes has primarily focused on three directions – 1. Driver gene identification based on mutational recurrence^5^; 2. Assessment of functional consequences of non-synonymous mutations; and 3. Discovery of mutational signatures^5–7^. Different from these, we asked if cancer related coding variants arise from different mutational processes as compared to germline or other non-cancer somatic mutations. This question is of significant clinical relevance since a key constraint in genomic testing in oncology is that matched normal tissue specimens are usually not collected. As a result, a large number of uncharacterized or not-so-well-characterized genomic alterations go unrecognized. The only visible effort in this direction is SGZ (somatic-germline-zygosity), a computational method for predicting somatic vs. germline origin of genomic alterations^8^. The authors of the study used ultra-deep (> 500x) targeted sequencing of 394 cancer-related genes for making confident somatic variant calls. To boost the prediction accuracy further, the authors factored in information about tumor content, ploidy, and copy number. We asked if ASSM based learning can discern cancer-specific SNVs based on sequence information alone. As indicated above, we restricted our analysis to the Chromosome X. We posed the problem of detecting cancer mutations as a supervised classification task. For Chromosome X, a total of 34,981/66,165 unique SNVs were obtained from ExAC/COSMIC databases^1, 9^, respectively, to prepare the training data for the task of predicting germline/non-cancer somatic vs. somatic origin of genomic alterations. These numbers were inflated to 285,102/590,171 upon considering the splice variants of the associated 332 protein-coding genes. Notably, the Chromosome X feature about 800 protein-coding genes. Reduction in the number of genes was due to relatively low alteration counts or extremity of their lengths (Fig. 1B-F). Since synonymous alterations are expected to have the least effect on cellular fitness, we ignored them in our downstream analysis.

To construct the training and the validation sets, as a stricter mechanism, we split the alterations based on genes of origin. In this way, SNVs sourced by 98 out of the 332 genes were held out for validation (Fig. 1D). We constructed a neural network architecture for classifying Aminoacid Switch Sequences (Fig. 2A), wherein 300 sized vectors (as discussed above) were used to represent the individual switches. The network architecture consists of a non-trainable layer for pre-trained embeddings followed by two *bi-directional Long Short-Term Memory* (bi-LSTM)^10, 11^ layers. These layers are then followed by a *time-distributed* layer and *batch normalization* layer. Time-distributed layer weights are shared across all the time-states along the switch sequences. Of note, the time states do not communicate with each other. Further, we implanted an *attention* layer^12^, followed by a *dense* layer with *sigmoid* output to classify the sequences. *Binary cross-entropy* loss was minimized to train the network ^13^. The proposed network architecture uses a total of 3,877,201 parameters, of which 194,600 are non-trainable. We obtained an Average Precision (AP, i.e., the area under the precision-recall curve) of 0.75, indicating classifiability of the mutational subtypes (Fig. 2B). For the control group, we constructed two fake classes of alterations by randomly splitting the pool of SNVs obtained from the ExAC browser. As expected, we obtained an AP of 0.5. With COSMIC alterations, our finding was similar (Fig. 2B). This tends to strongly support the deduction that cancer specific SNVs fall into a very different distribution, and are not uniformly distributed.

**Figure 2.**
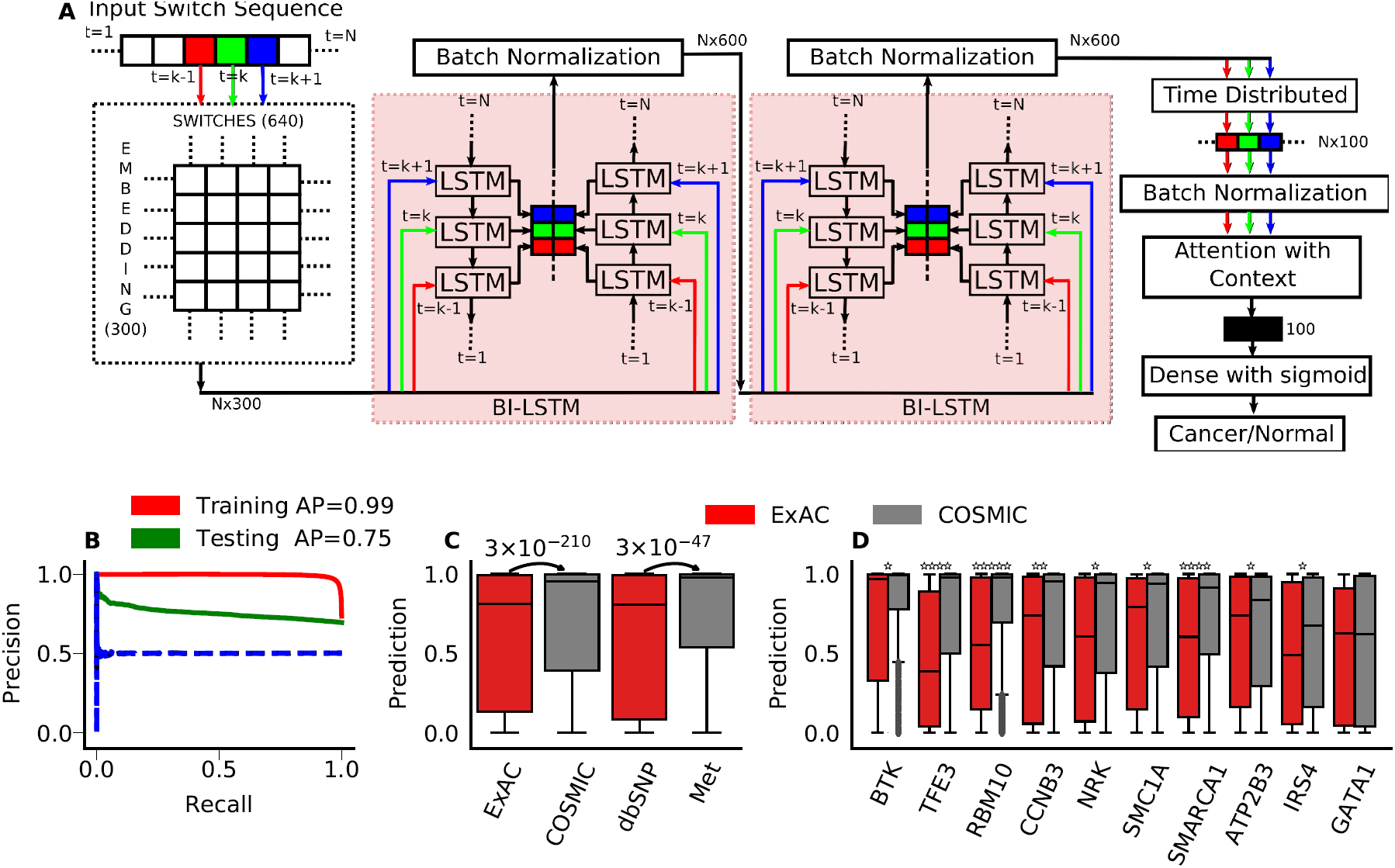
(*A*) The deep learning architecture used for training Aminoacid Switch Sequences constructed using alterations obtained from ExAC and COSMIC. (*B*) Precision-Recall (PR) curve obtained after 200 epochs. Red and green curves show performance on the training set and the validation set, respectively. Training and validation performances measured on fake alteration classes, constructed by randomly splitting cancer/non-cancer alterations into two equal size groups. All four PR curves thus obtained collapsed on the 0.5 Precision line. (*C*) Boxplots depict the distribution of the prediction scores (probability of being a cancer alteration) assigned to the ExAC and COSMIC alterations in the validation set. Similar trend can be observed for non-pathogenic dbSNP alterations and putative driver mutations reported by Priestley and colleagues^15^. (*D*) Boxplots show the distribution of the prediction scores assigned to ExAC and COSMIC alterations for the known driver genes from the validation set.

For an independent validation of the trained model, we obtained the neutral SNVs from the dbSNP database after removing the entries which are tagged pathogenic^14^. For a matching cancer alteration pool, we considered somatic SNVs (potential drivers) from a recently published pan-cancer study of solid metastatic tumors^15^ (referred to as *Met*). Notably, this study sequenced and analyzed 2,520 tumor samples representing the Dutch population. Following similar filtering criteria as discussed above, and removing sequences that overlapped with the training set, we were left with 315,040 non-cancer and 1,220 cancer-related mutations on Chromosome X. We predicted the cancer origin of the SNVs using the model, trained on ExAC and COSMIC data. As expected, cancer mutations were assigned relatively higher prediction scores (Mann-Whitney U-test *P*-value ¡ 3.01 × 10^−47^), thereby underscoring the robustness and cross-demographic reproducibility of our predictions (Fig. 2C).

Driver genes play a pivotal role in the diagnosis and clinical management of cancers. As such, we asked if our model differentiates between driver gene-specific germline (largely) and somatic mutations. Combining multiple driver gene databases (OncoKB^16^, intOGen^17^ and Cancer Genome Interpreter (CGI)^18^) offered us 55 potential drivers that belong to Chromosome X. Only 10 of these (RBM10, IRS4, CCNB3, ATP2B3, NRK, BTK, SMC1A, SMARCA1, GATA1, and TFE3) were not used at the time of training. We subjected both categories of the mutations to the predictor. Except for GATA1, for all other genes, we observed significant differences in the distribution of the prediction scores (Fig. 2D).

In this report, we put forth a mechanism to learn the embedding of coding variants by digesting a large pool of exome sequencing data. We demonstrated its utility in detecting cancer-related somatic mutations without the need for matched normal tissue samples. In other words, our findings suggest that cancer-specific SNVs, including the passenger mutations, occur with differential nucleotide context as compared to coding variants observed in healthy populations. One significant advantage of ASSM is that it is not reliant on any clinical or pathological parameters. We predict that the proposed approach could be adopted in attempting a broad range of questions concerning genotype-phenotype interlinking. Some potential use-cases include causative gene inference in rare diseases, assessing the impact of mutations on protein conformation, and exploring the role of splicing mutations in genetic disorders.

## Acknowledgements

We thank Gaurav Ahuja, Giovanni Ciriello, and Kedar Natarajan for their valuable comments and suggestions. This work is partially supported by the INSPIRE Faculty Grant [DST/INSPIRE/04/2015/003068] awarded to DS by the Department of Science and Technology (DST), Govt. of India.

## Author Contributions

D.S. conceived the study. D.S. and J. designed the experiments. P.G. and A.J. wrote the code and performed all the analyses under the supervision of D.S. and J. All the authors discussed the results, co-wrote, and reviewed the manuscript.

